# Supported membrane assay probes PLCγ1 activity in LAT condensates

**DOI:** 10.64898/2026.01.23.701188

**Authors:** Joseph B. DeGrandchamp, Sameer V. Rajesh, L. J. Nugent Lew, Jay T. Groves

**Author notes:** Institute for Digital Molecular Analytics and Science, Nanyang Technological University, Singapore, Singapore (Joseph B. DeGrandchamp). UT Southwestern Medical School, Dallas, Texas, United States of America. Corresponding Author (JTG).

## Abstract

Phospholipase C-γ1 (PLCγ1) plays a critical role linking T cell receptor activation with downstream signaling pathways including calcium. PLCγ1 activation in T Cells relies on phosphotyrosine-mediated recruitment to the membrane-bound scaffold LAT, which becomes crosslinked through a bond percolation network with Grb2 and other scaffold and signaling molecules to form a signaling condensate. PLCγ1 in these LAT condensates becomes activated, leading to induction of extracellular calcium influx. While PLCγ1-driven calcium signaling is clearly correlated with LAT condensation, it is less clear how—or if— the LAT condensation state facilitates PLCγ1 activity. Here we develop an image-based PLCγ1 activity assay in supported bilayers that enables simultaneous measurement of both PLCγ1 recruitment to phosphorylated LAT and PLCγ1-catalyzed hydrolysis of PIP_2_ in the membrane. The condensation state of LAT is independently controlled by adjusting levels of co-condensation proteins such as Grb2, SOS, GADS, and SLP76. The hydrolysis product, diacylglycerol (DAG), remains in the membrane and is monitored as a readout of catalytic activity using a DAG sensor based on the C1b (DAG binding) domain of PKCθ. Assays are performed directly with mammalian cell lysate containing fluorescent PLCγ1 fusion constructs. The results reveal that PLCγ1 is highly active when recruited to dispersed LAT and that the condensed state does not promote activity. Overall, this assay platform reveals that despite the correlation between PLCγ1 signal gating and LAT condensation, the physical environment of the condensate itself is not a key regulator of PLCγ1 signaling. More broadly, this assay system offers a quantitative means of probing how PLCγ1 activity is controlled at the membrane.

## Introduction

T cells have the remarkable ability to detect a single foreign antigen molecule among many thousands of self-antigens on an antigen presenting cell surface [1–4]. This single molecule sensitivity places tremendous demands on both fidelity and signal amplification capabilities of the T cell receptor signaling pathway. Antigen discrimination originates from various events shaping full T cell activation [5,6], but most relevant to the present study is the condensation of phosphorylated LAT (pLAT) on the membrane [7–11].

LAT has four key tyrosine residues that are phosphorylated by the kinase Zap70 upon T cell receptor (TCR) activation [12]. Three of these (pY171, pY191, and pY226) bind the cytosolic adaptor Grb2 [12–14]. Through multivalent interactions with pLAT and the Ras-activator SOS, Grb2 can crosslink pLAT into extended assemblies that have been referred to as protein condensates [8,10,15,16]. In the case of SOS, autoinhibition release and initiation of Ras activation is strongly dependent on the condensation state of LAT, and is controlled by a distinct type of kinetic proofreading mechanism [15,16]. The fourth phosphorylation site on LAT, LAT-pY132, is selective for Phospholipase C-γ1 (PLCγ1) [12–14]. PLCγ1 becomes activated upon recruitment to pLAT and plays a key role initiating signaling steps leading to extracellular calcium influx. This calcium signal is among the earliest cell-wide signals that show distinct single molecule sensitivity. Recent observations have revealed that individual TCR activation events produce discrete LAT condensates and that each condensate contributes a single calcium spike to the developing cell-wide calcium signal [11,17,18]. A quantitative impulse-response function for LAT condensation to calcium has been measured in this previous work. The results indicate almost immediate (<10 s) activation of the full extracellular calcium flux after initial condensation of each LAT condensate, followed by a rapid attenuation process without oscillation or ringing. This suggests a remarkable level of control over PLCγ1 activity from the condensate, and it is tempting to hypothesize that PLCγ1 autoinhibition release may exhibit similar kinetic regulation as observed for SOS. However, there is limited experimental support for this idea and, as will become clear from the results described below, skepticism is warranted.

PLCγ1 is a large, multi-domain enzyme that hydrolyzes phosphatidylinositol-4,5-bisphosphate (PIP_2_) at the cell membrane into the second-messengers diacylglycerol (DAG) and inositol-triphosphate (IP_3_) [19]. In T cells, DAG feeds into the MAPK pathway through Ras, while IP_3_-triggered calcium release is linked to transcription of the gene for IL-2, an inflammatory cytokine that marks T cell activation [6]. PLCγ1 is strictly regulated by an autoinhibitory loop that obstructs its active site and membrane binding domains [19]. This loop contains, in sequence, an nSH2 domain, a cSH2 domain, and an SH3 domain. The nSH2 of its autoinhibitory loop binds strongly and specifically to pY132 on pLAT, while the loop’s SH3 domain additionally links PLCγ1 to pLAT by binding the cytosolic scaffold SLP76. SLP76 binds to the SH3 domain of the Grb2-like adaptor Gads, which in turn can bind back to pLAT at either pY171 or pY191 (competing with Grb2). Ultimately, this results in a multi-protein complex (PLCγ1:SLP76:Gads:LAT) facilitating the interaction of PLCγ1 with LAT and the membrane, where PLCγ1 can be phosphorylated at Y783 by ITK (also recruited to SLP76) [6,19–22]. Upon phosphorylation, the autoinhibitory loop’s cSH2 domain binds intramolecularly to pY783, leading to conformational rearrangements that activate the enzyme by releasing the active site for substrate binding.

PLCγ1 has been observed to crosslink pLAT as well, further tying the LAT condensate to PLCγ1 and immediately calling into question the effect of pLAT condensation on PLCγ1 activation and enzymatic turnover—and thus T Cell activation itself [23]. While PLCγ1 activation has been studied using purified constructs and solution assays [19,21], a technique to simultaneously quantitate both its recruitment to pLAT and catalytic activity in a membrane context where LAT can condense has not existed. We previously developed a supported lipid bilayer (SLB) method to study active full-length SOS from mammalian cell lysate in LAT condensates [15,24]. Herein, we apply a similar strategy to PLCγ1 using PLCγ1-mNeonGreen (mNG) fusion constructs from lysate flowed over supported bilayers functionalized with pLAT and the PLCγ1 substrate PIP_2_. PLCγ1 activity is tracked by monitoring DAG build up in the membrane using a fluorescently labeled DAG binding domain (C1b domain of PKCθ) [25,26]. Using TIRF imaging, we determine the spatial organization of analytes at the membrane, concurrently observing PLCγ1 activity and pLAT condensation states. The results confirm the findings of previous studies showing the central importance of pLAT recruitment and Y783 phosphorylation to enzymatic activation. However, we also find that PLCγ1 is highly active when recruited to dispersed pLAT and that LAT condensation provides no activity enhancement. PLCγ1 regulation by LAT condensation clearly utilizes a control mechanism drastically different from SOS, even though they are both signaling from the same condensate. More broadly, this assay presents a useful tool for further reconstitution of PLCγ1 activation dynamics and mutational analyses to determine mechanisms of PLCγ1 signaling and its regulation by condensates on the membrane.

## Results

### Quantitative activity assay for PLCγ1 from cell lysate

PLCγ1-mNG was transfected into HEK293T cells to facilitate activation studies of the full-length protein from lysate. Lysate from the cells (generated by minimal sonication) was clarified by benchtop centrifugation, diluted in an imaging buffer (with phosphatase inhibitors), and flowed over pLAT/Hck-functionalized supported lipids bilayers (SLBs) with 2 mol% PIP_2_ content (Fig 1A). Phosphorylation of LAT is ensured by co-functionalization of Hck kinase to the membrane according to established methods [10,15], which we later determine is also capable of phosphorylating PLCγ1-mNG at the membrane. PLCγ1-mNG recruitment to LAT was measured via TIRF imaging of surface mNG signal (Fig 1C). To determine whether PLCγ1-mNG actively hydrolyzed PIP_2_ to DAG, a DAG sensor was developed based on the C1b (DAG binding) domain of PKCθ fused with an N-terminal SNAP-tag (Alexa 647 labeled) (Fig 1A). A similar construct has been utilized in live cells, and other labeled binding domains have been well-characterized for use in reactions on SLBs [15,25,26]. Surface fluorescence of the DAG sensor was tracked in sequence with PLCγ1-mNG (Fig 1C), with a low background signal giving way to a marked increase in intensity in line with PLCγ1-mNG recruitment.

**Fig 1.**
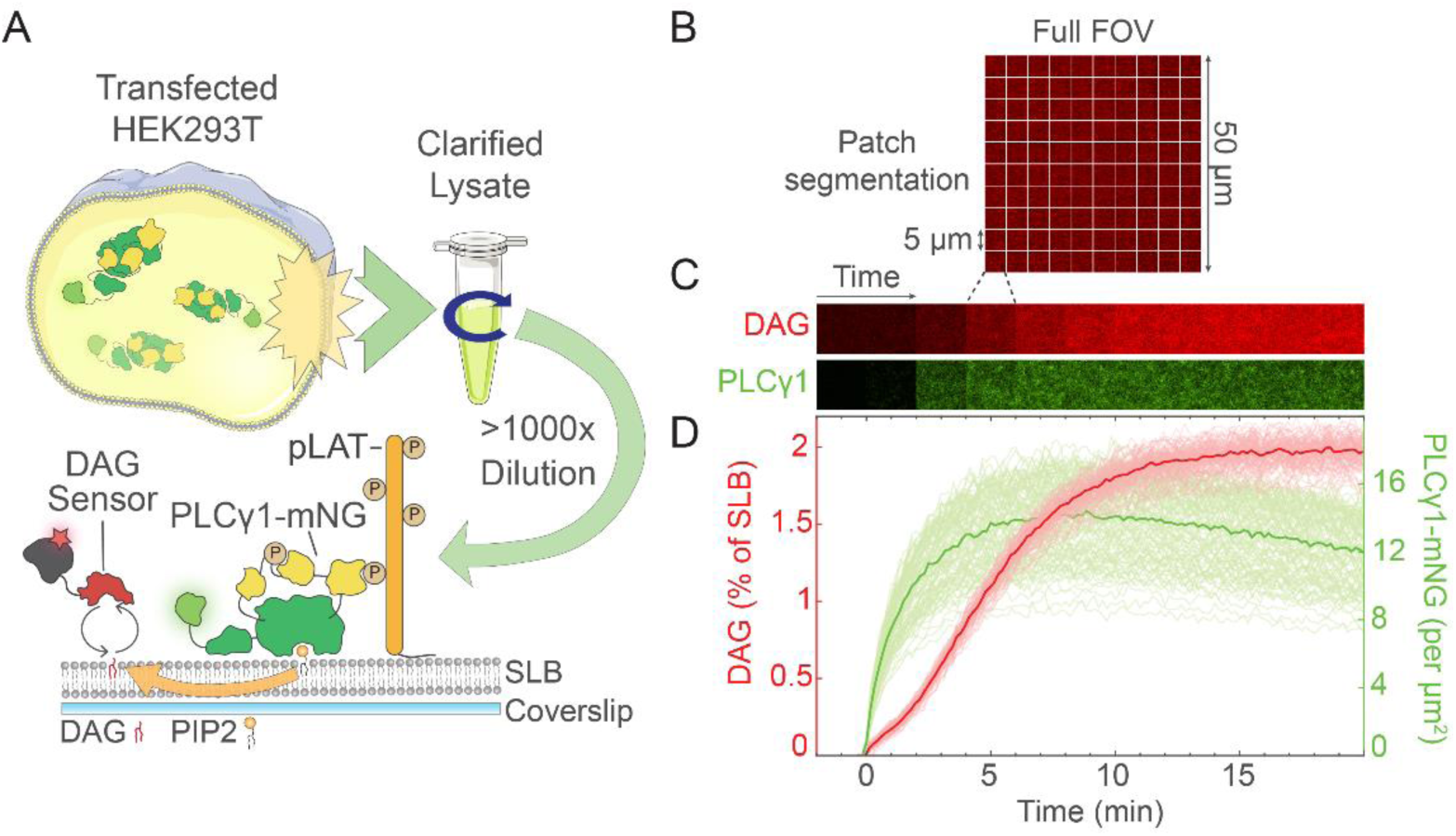
PLCγ1 dynamics are measured directly from lysate by TIRF imaging of supported lipid bilayers. (A) A schematic summarizing the lysate assay protocol. PLCγ1-mNG was generated in HEK293T cells, then flowed directly as dilute lysate over pLAT bilayers containing PIP_2_. Active PLCγ1-mNG hydrolyzed PIP_2_ to DAG, which was detected using a labeled DAG sensor (PKCθ-C1b-SNAP-Alexa 647). Parts of the figure were drawn using images from Servier Medical Art Commons Attribution 3.0 Unported License (http://smart.servier.com (accessed on 21 March 2022)). (B) An example TIRF image of the DAG sensor demonstrating the segmentation of 50 x 50 µm images for analysis. TIRF images in a time series were sectioned into 5 x 5 µm patches and analyzed as separate instances. (C) Images of a single 5 x 5 µm patch in the DAG sensor channel (Top) and PLCγ1-mNG channel (Bottom) through time, corresponding to the time axis for the curves below. (D) Curves obtained from processing the raw median fluorescence from membrane patches, representing the amount of DAG in the bilayer (red, left axis, foreground curves) and density of PLCγ1-mNG recruited (green, right axis, background curves) over 20 minutes. PLCγ1-mNG lysate (∼1:1000 dilution) with DAG sensor was injected at *t =* 0 min. To attain quantitative curves, raw fluorescence was first normalized to the time of injection. DAG was calibrated using the assumption that the average normalized maximum for multiple experiments must be 2% (given 2% PIP_2_ bilayers). PLCγ1-mNG was calibrated by matching single-molecule counts (at high laser power) to bulk fluorescence (at low laser power). The two solid, darkly colored curves represent the median of all patches analyzed. Lightly colored curves in the background represent the results from each single patch (n = 100). Lipid composition (mol%): 94:4:2 DOPC:Ni-DGS:PIP_2_. LAT density: ∼1500 µm^-2^.

The use of supported membrane corral arrays, partitioned by grids of physical barriers micropatterned onto the underlying substrate, for the analysis of signaling reactions has been well-established [15,27–30]. Herein, we utilize a similar but distinct strategy by software segmentation of bilayers that are physically unpartitioned. TIRF images with a full Field of View (FOV) of 2500 µm^2^ were separated into 25 µm^2^ patches and analyzed separately (Fig 1B). This serves both to correct differences across the image due to uneven illumination and aberrations and to reduce the effects of spurious outliers (such as rare aggregates or membrane defects) which are often uncharacteristically intense. Median fluorescence across each patch was computed at each timepoint to yield raw fluorescence curves. For quantification, a few reasonable assumptions were applied. Curves were first normalized to the point of injection. With the maximum mol% of DAG in the SLB fixed at 2% (capped by the amount of PIP_2_ in the bilayer), it can be assumed the eventual plateau of normalized DAG fluorescence signal corresponds to 2 mol% of DAG. The relative increase and plateau of normalized DAG signal were verified to be consistent over several experiments and over multiple days, allowing for the calculation of an average “intensity to DAG” coefficient to be applied across varied conditions (described in detail in the *Materials and Methods*). PLCγ1-mNG signal was similarly normalized and converted to PLCγ1-mNG densities at the bilayer by correlating bulk images (at low laser power) to images of countable PLCγ1-mNG (at high laser power) for varying PLCγ1-mNG concentrations. The resulting quantitative curves for a typical PLCγ1-mNG lysate assay on a pLAT bilayer show an increase in DAG following shortly after PLCγ1-mNG recruitment and an eventual total conversion of PIP_2_ after roughly 20 minutes (Fig 1D). There is good agreement across all patches analyzed (shown as lightly colored curves in the background). PLCγ1-mNG signal starts to decrease after roughly 90% of PIP_2_ has been hydrolyzed, potentially due to decreased membrane binding in the absence of substrate (in experiments with labeled LAT, there was no appreciable desorption of LAT detected over the experiment timeframe). The DAG production curves appear potentially sigmoidal, with a differential rate noticeable in the initial few minutes. However, in the assay’s current form, this is hard to separate from solution perturbation introduced by the injection of the lysate/sensor mixture into the channel and is not interpreted.

### PLCγ1 activity requires phosphorylation and pLAT recruitment

A western blot of control cell lysates and lysates containing PLCγ1-mNG probed for total PLCγ1 content indicated the presence of endogenous PLCγ1 (Fig 2A). However, exogenously expressed PLCγ1-mNG was roughly six times more abundant relative to the endogenous species (Fig 2B), in line with the ratio found for past results for SOS-EGFP from lysate [24]. In addition, PLCγ1-mNG showed good stability over 60 minutes at room temperature, with limited changes in lower molecular weight bands indicative of proteolysis (Fig 2A).

**Fig 2.**
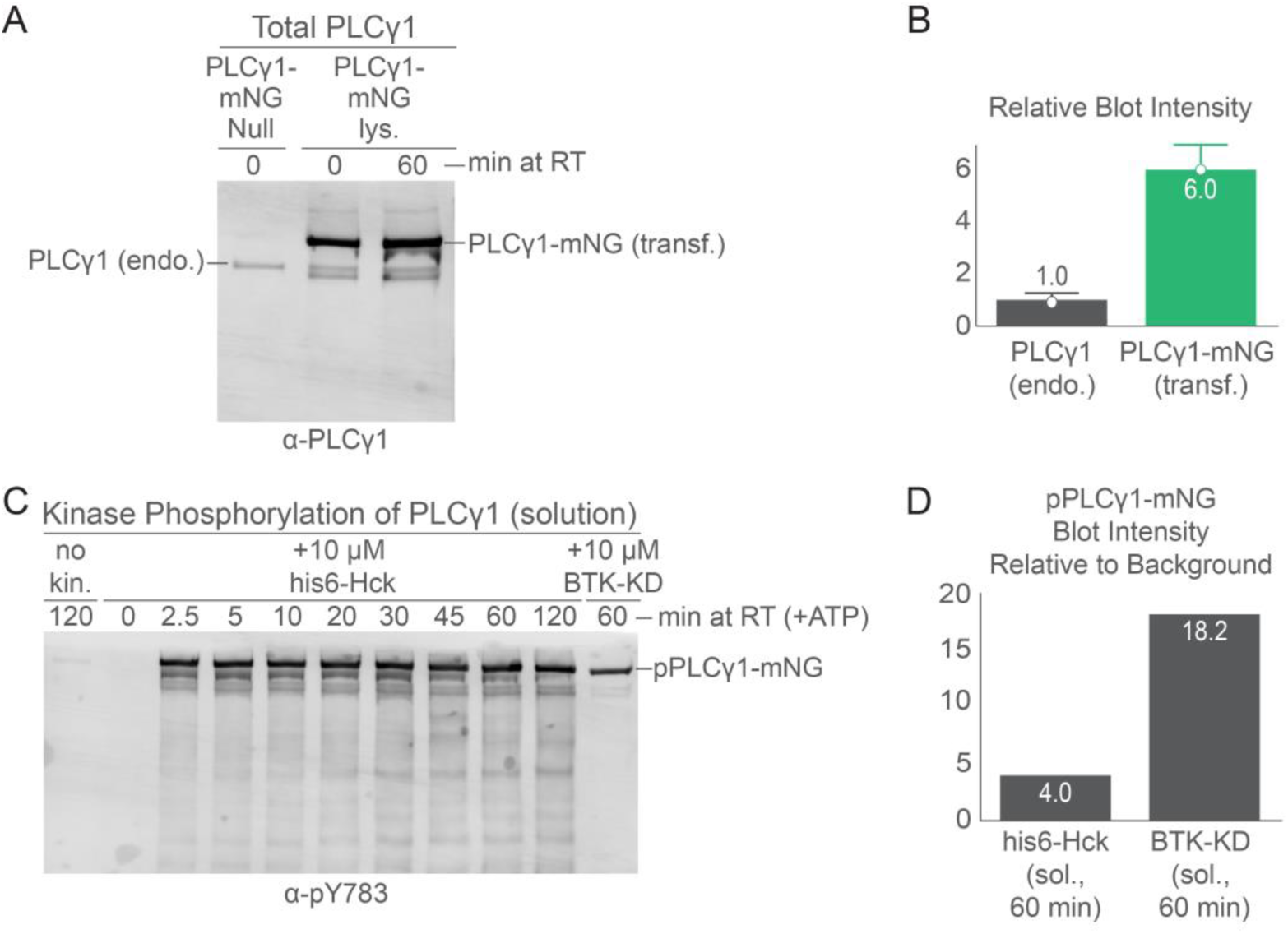
Exogenous PLCγ1-mNG is more abundant than endogenous species and activatable by *in vitro* phosphorylation. (A) Western blot probed for total PLCγ1 content. Cells were transfected with either a Gads expression vector (“PLCγ1-mNG Null”, left lane) or PLCγ1-mNG vector (middle and rightmost lanes). PLCγ1-mNG lysates were loaded after either 0 min (middle) or 60 min (right) at RT to determine the stability of PLCγ1-mNG against degradation. Each lane was loaded with 1 µL of lysate. (B) Relative blot intensities (background corrected) for endogenous PLCγ1 (left, grey) versus transfected PLCγ1-mNG (right, green) normalized to endogenous levels (same lane) (n=2). (C) Western blot probed for pY783 with a series of time points and different kinase conditions. Phosphorylation reactions were conducted in 10 µL volumes containing 1 µL of PLCγ1-mNG lysate each, ATP, phosphatase inhibitor, and kinase in buffer. Reactions were kept at RT for the designated time, then quenched with SDS-PAGE loading buffer. Either no kinase, his6-Hck, or BTK-KD (Kinase Domain) was used. (D) Blot intensities (background corrected) of pPLCγ1-mNG relative to non-specific pY bands.

While there is a visible band (below endogenous PLCγ1) that may correspond to the absence of mNG (due to cleavage or misfolding), the majority of PLCγ1-mNG remained intact. The small difference in the lower band may be due to species differences. PLCγ1-mNG was constructed using the murine sequence due to availability, while the endogenous species in HEK293T cells would have the human sequence. Importantly, the two sequences have 97% residue similarity and a difference of only 12 residues in length (∼1 kDa), and the differences are not expected to affect results strongly as evidenced by the frequent use of rat PLCγ1 in other studies [19–21].

With regards to the activation of PLCγ1 by phosphorylation, a pY783 specific antibody reveals PLCγ1 can be strongly phosphorylated *in vitro* (Fig 2C). There is essentially zero phosphorylation of PLCγ1 at an initial time point (aliquots were thawed after being flash-frozen immediately following lysis). After two hours, there remained minimal detectable pPLCγ1, indicating a negligible contribution from endogenous kinases. Meanwhile, Hck, a commonly employed and abundant promiscuous kinase, phosphorylated PLCγ1 fully within a few minutes (Fig 2C) [10,15]. A dark band corresponding to pPLCγ1-mNG appeared quickly with Hck incubation and did not get more intense in subsequent time points, indicating maximal phosphorylation. The His6-Hck construct in this assay is also utilized on our SLBs to phosphorylate LAT (Fig 3A) and is thus the source of PLCγ1 phosphorylation in our assays. In addition, the kinase domain of BTK (BTK-KD) was able to phosphorylate PLCγ1 to a similar level in solution (Fig 2C). BTK activates PLCγ2 in B-Cells and is a close family member of ITK, the PLCγ1 activator in T-Cells. BTK exhibited remarkable selectivity for the phosphorylation of PLCγ1 compared to Hck (Fig 2D, S1 Fig).

**Fig 3.**
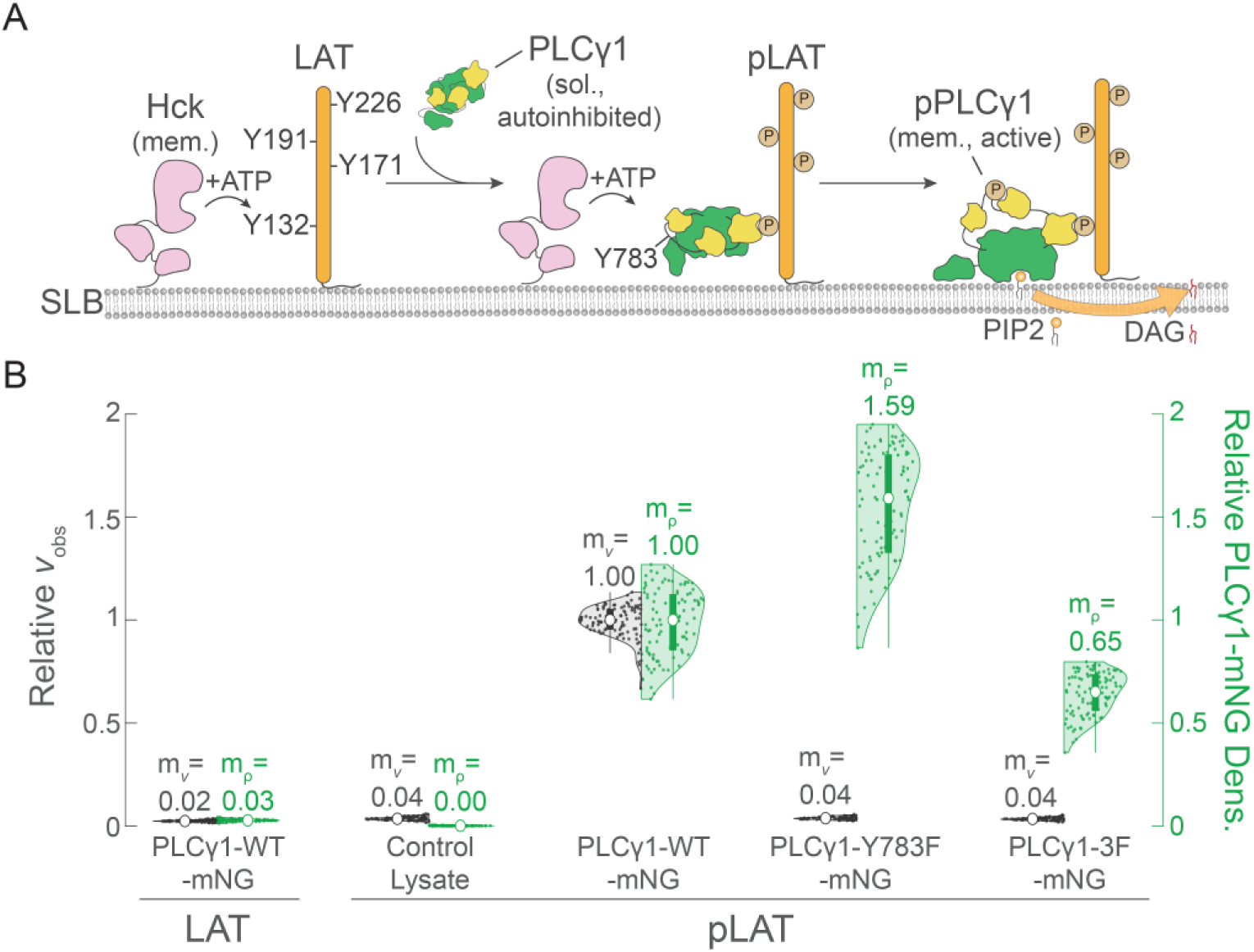
PLCγ1 requires pLAT and phosphorylation at Y783 to activate on the membrane. (A) A schematic showing greater detail of the lysate experiment at the membrane. Hck and LAT are functionalized to the SLB via Ni-histidine chelation. Hck phosphorylates LAT allowing for the recruitment of PLCγ1-mNG. PLCγ1-mNG is then subsequently phosphorylated by Hck at Y783 and is active at the membrane after structural rearrangement. (B) Relative observed enzymatic velocities (*v*_obs_) (black, left axis, defined as the enzymatic rate for the linear portion of the reaction trace) and PLCγ1-mNG densities (green, right axis, defined as density at the midpoint of *v*_obs_ measurement) for various assay conditions, normalized to the median value for PLCγ1-WT-mNG lysate on a pLAT bilayer. Distributions of 100 membrane patches are shown as half-violin plots, with single observations shown as points and distributions represented by a shaded area. Compact box plots are overlayed over the distributions to represent the median (white circles), quartile regions (box), and extremes with outliers excluded (whiskers). Median values for *v*_obs_ (m*_v_*) and PLCγ1-mNG density (m_ρ_) are expressly denoted for clarity. Lipid composition (mol%): 94:4:2 DOPC:Ni-DGS:PIP_2_. LAT densities: ∼1500 µm^-2^.

To further characterize the current assay and the requirements for PLCγ1 activation, a careful series of control conditions and PLCγ1 mutations were examined (Fig 3B). To facilitate comparisons between conditions, an observed enzymatic velocity (*v*_obs_) was extracted from the linear portion of each DAG production curve. For example, the slope of the previously discussed DAG curves was calculated between 2.5 and 5 minutes (Fig 1D). To simplify comparisons of PLCγ1-mNG density, the midpoint of the *v*_obs_-measurement window for each patch was considered. *v*_obs_ and PLCγ1-mNG density values were tabulated for each membrane patch and each condition, then normalized relative to the median of the PLCγ1-mNG-WT condition and visualized using dual-axes half-violin plots (Fig 3B). In the absence of pLAT (no Hck on the membrane), PLCγ1 recruitment and activity were both abolished. Control lysate showed a similar lack of activity, with a median *v*_obs_ (m*_v_*) corresponding to only 4% of the m*_v_* for PLCγ1-mNG-WT lysate (and no PLCγ1-mNG, as would be expected).

To assess the importance of Y783 to activation, phenylalanine mutants were generated to abrogate PLCγ1 phosphorylation. We introduced the Y783F mutation as well as a triple mutation, referred to as “3F,” that included Y771F and Y775F (adjacent PLCγ1 tyrosines also implicated in *in vivo* activation) in addition to Y783F [31]. Lysate from cells transfected with these constructs showed robust recruitment of PLCγ1-mNG to pLAT but had corresponding *v*_obs_ equivalent to control lysate (Fig 3B). Our results show the Y783F mutation was sufficient to fully inhibit PLCγ1. Differences in observed PLCγ1-mNG density between the WT and mutants were most likely due to expression levels of the constructs in the HEK293T cells rather than anything mechanistic. While transfection efficiencies were roughly 80% for all constructs and the numbers of cells lysed were similar, cell-to-cell or cell cycle variation in expression was not accounted for. PLCγ1-mNG-WT also showed similar batch-to-batch differences (a single batch of cell lysate was used for all experiments in this study). Closer examination would be required to fully decipher any differences in recruitment between the constructs.

These results are in agreement with previous studies claiming PLCγ1 requires both a phosphorylated receptor and phosphorylation of pY783 for full activation at the membrane [19,21]. Notably, even pPLCγ1-mNG (pre-phosphorylated in solution by BTK-KD) does not recruit to or hydrolyze PIP_2_ in membranes devoid of LAT (S2B Fig). The full mechanism of our *in vitro* lysate assay is then most likely to be as depicted in Fig 3A. Hck first phosphorylates LAT on the bilayer, PLCγ1 from lysate is introduced and recruits to pLAT, PLCγ1-Y783 is then also phosphorylated by Hck, and ultimately PLCγ1 actively hydrolyzes PIP_2_ to DAG. With the basics of the assay established, additional components from the *in vivo* signaling system were added to illuminate further PLCγ1 interactions.

### PLCγ1 localizes strongly to pLAT condensates but displays diminished activity

Recent studies of PLCγ1 in a condensate-forming growth-receptor pathway suggest condensation plays a role in increasing the observed activity of PLCγ1 (23). pLAT has been shown to form condensates primarily via crosslinking by Grb2 and SOS in cells and *in vitro*, and PLCγ1 was recently seen to participate in these condensates [23]. Therefore, it seemed likely pLAT condensation would have an effect on PLCγ1 activity. To elucidate any possible effect, we utilized purified forms of Grb2 and SOS-PR to condense LAT while performing our PLCγ1 lysate assay. With the addition of these adaptors, labeled LAT partitioned into strongly condensed phases as expected, with no visible effect from the presence of lysate (Fig 4A). PLCγ1-mNG also localized into these condensed phases, as seen in TIRF images where shapes formed by PLCγ1-mNG are similar to those of LAT-Alexa 555 (Ax555) (Fig 4A-B). The DAG sensor was unaffected at the imaging time utilized and remained visibly homogeneous (Fig 4C).

**Fig 4.**
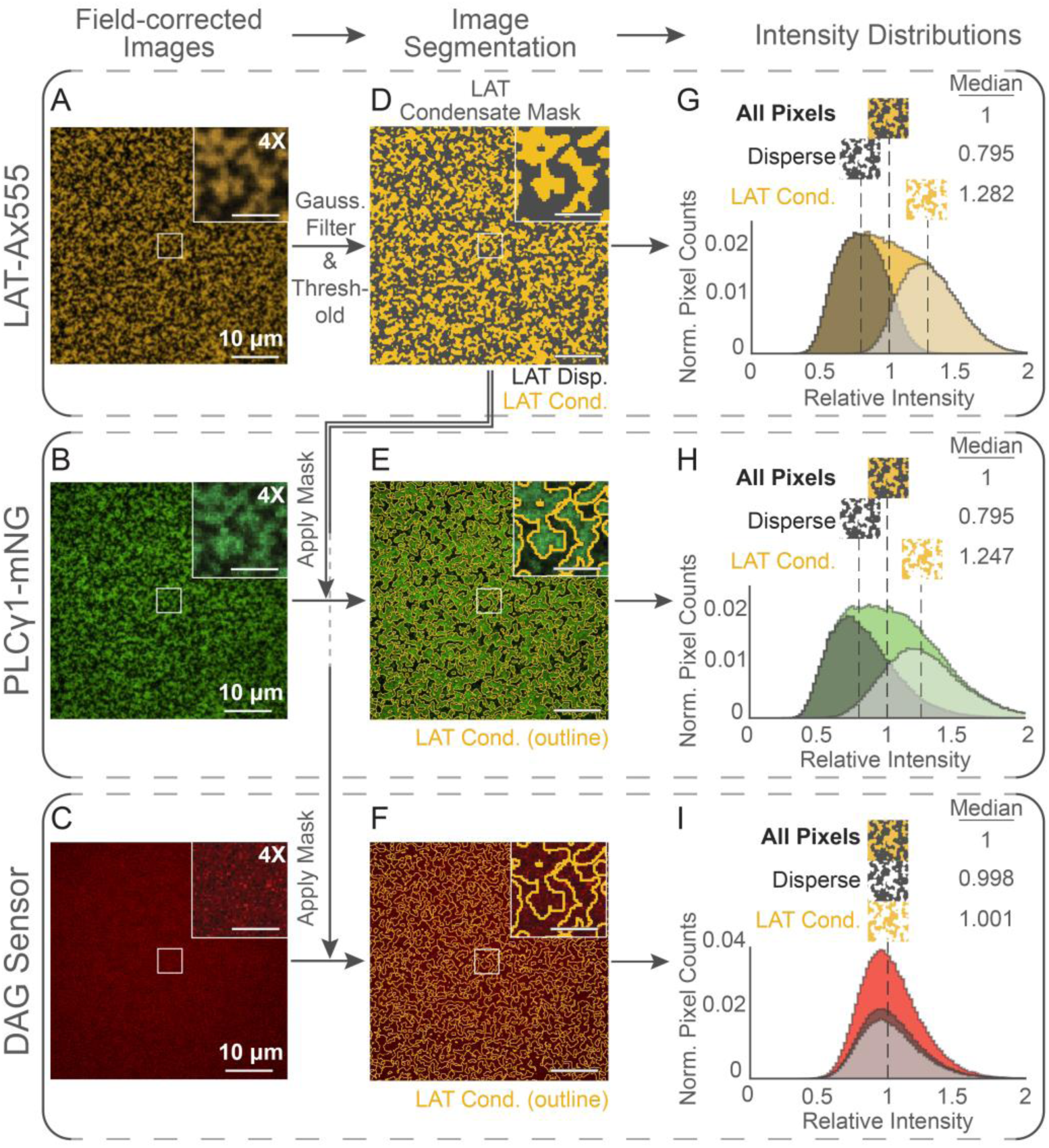
PLCγ1 localizes to the pLAT condensate, but DAG sensor is not affected at relevant timescales. (A)-(C) Field-corrected images of LAT-Alexa 555 (Ax555) (gold, top), PLCγ1-mNG (green, middle), and DAG sensor (red, bottom) with 5 µM Grb2 and 2 µM SOS-PR in solution at *t* = 15 min. The entire FOV is 50 x 50 µm (scale bar = 10 µm) with an inset showing a 4X expansion of the center 5 x 5 µm patch (scale bar = 2.5 µm). D. A binary image mask for LAT segmented into condensed (gold) and disperse (grey) areas obtained from Gaussian filtration (σ=1) and thresholding of the LAT-Ax555 image in (A). Inset and scale bars are equivalent to those in (A). (E)-(F) Images showing the effect of applying the binary LAT mask to PLCγ1-mNG (green) and DAG sensor (red) channels. Only the outline of the applied mask is shown for visibility. Insets and scale bars are equivalent to those in (B-C). (G)-(I) Histograms of relative pixel intensities for LAT-Ax555 (gold, top), PLCγ1-mNG (green, middle), and DAG sensor (red, bottom). Pixel intensities were normalized to the median value for all pixels. Values greater than 1 indicate an increase in brightness from the overall median, while values less than 1 indicate a decrease. Histograms for all pixels are displayed in full color in the background. Histograms for pixels that are in areas of condensed LAT are shown in translucent white in the foreground. Pixel histograms for disperse LAT areas are shown in translucent grey. Dotted lines denoting the median of each histogram are decorated with an icon corresponding to the LAT mask for all pixels (gold and grey), LAT condensed (gold only), or LAT disperse (grey only). Median fold change values for each histogram are displayed in the top right for clarity. Lipid compositions for all experiments (mol%): 94:4:2 DOPC:Ni-DGS:PIP_2_. LAT density: ∼1500 µm^-2^ (10% labeled).

To quantitatively characterize the localization of PLCγ1-mNG and determine any possible effects of the condensate on our assay, we employed an image analysis technique comprised of image segmentation and pixel mapping. We created a binary “LAT condensation mask” (0 for pixels outside condensed areas, 1 for pixels in condensed areas) by putting an image of LAT-Ax555 through a gaussian filter, then applying an appropriate threshold (Fig 4D). We then applied this mask to pixels from the original LAT image as well as images from the PLCγ1-mNG and DAG sensor channels (Fig 4E-F). The outline of the applied mask in Fig 4E contains most of the visible PLCγ1-mNG, while in Fig 4F there appears to be no difference in DAG sensor intensity inside and outside the masked area. Pixel intensities were normalized to the median of the entire image and then visualized in histograms (Fig 4G-I). Histograms for pixels across the entire image (unsegmented) for LAT and PLCγ1 show characteristics of multiple underlying (but unresolved) gaussian distributions (Fig 4G-H). However, by mapping pixels by the condensed/disperse state of LAT, the underlying distributions were successfully resolved. Pixels in LAT-condensed areas for both LAT-Ax555 and PLCγ1-mNG images had a median intensity 1.2-1.3 times brighter than the whole image (and ∼1.6 times brighter than disperse areas) (Fig 4G-H). Pixels in the corresponding DAG image were homogenously distributed, regardless of the mapped state of LAT, with differences in relative median intensity of less than 0.002 (Fig 4I). This substantiates the claim that PLCγ1 is detectably recruited to pLAT condensates, and that our DAG sensor is not affected by the dense matrix (at relevant timescales). The DAG sensor also eventually reaches the same saturation (of 2 mol%) as uncondensed conditions (S3A Fig). With this established, we could move on to analyzing the activity of PLCγ1 within condensates.

Interestingly, it is immediately apparent from measurements of *v*_obs_ in condensing conditions (with Grb2 and SOS-PR) that the activity of PLCγ1 is *decreased* in LAT condensates, even with an increase in overall PLCγ1-mNG density (Fig 5B). The median value for *v*_obs_ (m*_v_*) for the Grb2:SOS-PR condition is only 40% of the uncondensed condition, despite a nearly threefold increase in PLCγ1-mNG (maximum over 20 minutes). But while Grb2 and SOS are thought to be the main mediators of pLAT condensation and are capable of interacting with PLCγ1, Gads and SLP76 are the canonical partners of PLCγ1 in this system [20–22]. To determine potential differences in these adapters, Gads and SLP76 were expressed in HEK293T cells to generate lysate with full-length versions of these proteins (which have been shown to be troublesome in purified form) (S3B Fig) [20]. To make fair comparisons, control lysate was added to the other conditions to provide roughly equivalent amounts of endogenous protein among all trials. Activity due to endogenous PLCγ1 was higher than previous experiments but still reasonably low compared to PLCγ1-mNG conditions (S3C Fig). When Gads:SLP76 were added to PLCγ1 alone, they had little effect on recruitment or activity (within error) and did not lead to condensate formation (Fig 5A-B). However, when added in addition to Grb2:SOS-PR in condensing conditions, m*_v_* nearly doubled and peak PLCγ1-mNG density returned closer to uncondensed levels. The structure of the condensation also appeared changed (Fig 5A).

**Fig 5.**
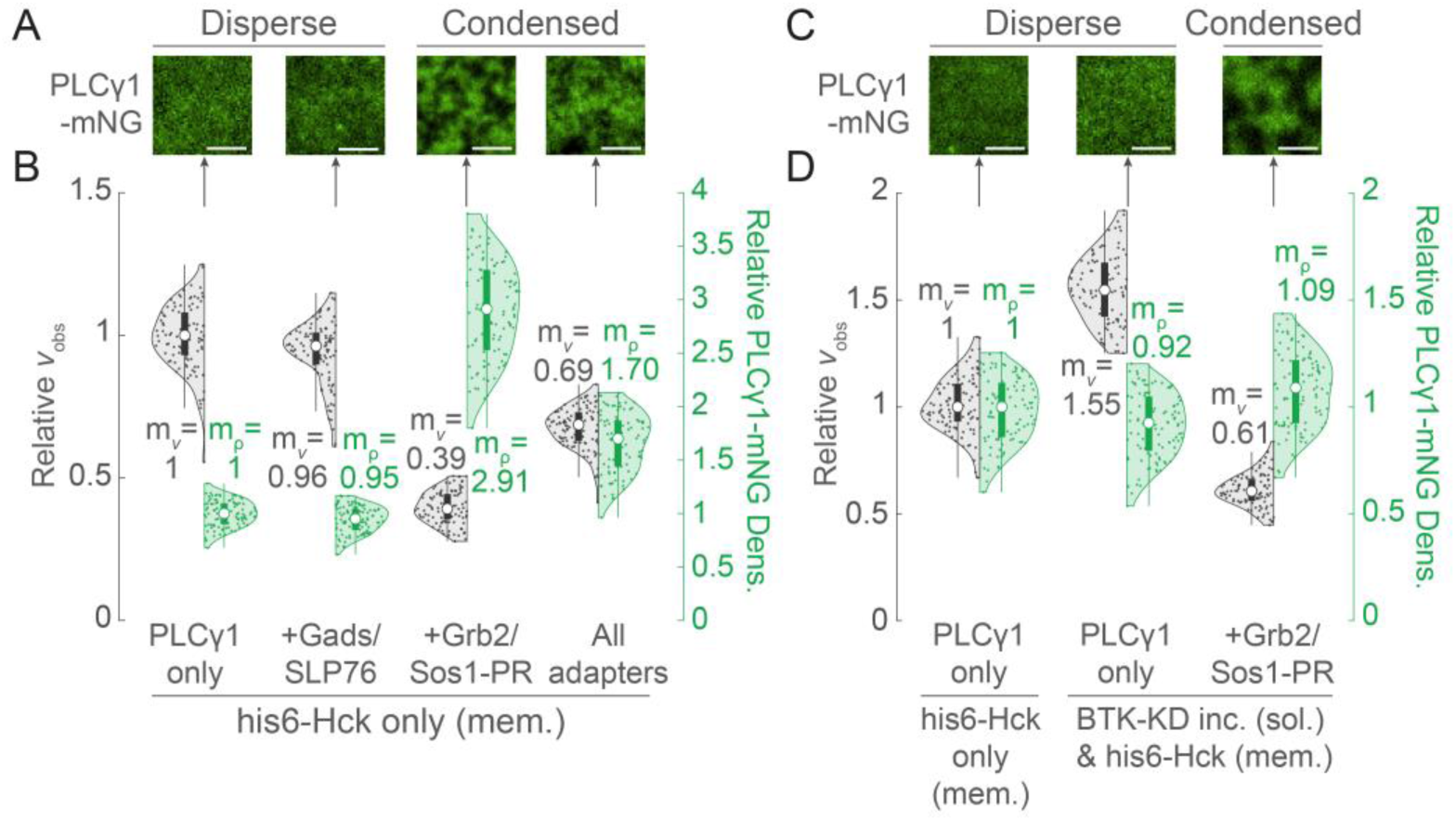
PLCγ1 activity is reduced in LAT condensates, regardless of phosphorylation state. (A) TIRF images (5 x 5 µm, scale bars = 2 µm) of PLCγ1-mNG in different conditions corresponding to the distributions below. From left to right: PLCγ1-mNG only (with 2 µL control lysate), PLCγ1-mNG + Gads:SLP76 (from lysate, 1 µL each), PLCγ1-mNG + Grb2:SOS-PR (purified, 6 µM:2 µM respectively), and PLCγ1-mNG + all previously described adapters. LAT-Ax555 (10% labeled) was used in this experiment but was determined to have negligible contribution in the PLCγ1-mNG channel. (B) Half-violin plots showing distributions of relative *v*_obs_ (grey, left axis) and relative PLCγ1-mNG density (green, right axis), normalized to PLCγ1 only conditions. Compact box plots overlay the distributions to convey statistics. Median values for *v*_obs_ (m*_v_*) and PLCγ1-mNG density (m_PLCγ1_) are expressly denoted for clarity. (C) TIRF images (5 x 5 µm, scale bars = 2 µm) of PLCγ1-mNG in different conditions corresponding to the distributions below. From left to right: PLCγ1-mNG only (no control lysate), PLCγ1-mNG pre-phosphorylated by 1 µM BTK-KD in solution for 1 hr, and similarly pre-phosphorylated PLCγ1-mNG + Grb2:SOS-PR (purified, 6 µM:2 µM respectively). Hck was on the membrane for all experiments. Unlabeled LAT was used in this experiment. (D) Half-violin plots showing distributions of relative *v*_obs_ (grey, left axis) and relative PLCγ1-mNG density (green, right axis), normalized to PLCγ1 only conditions (no pre-phosphorylation). Compact box plots overlay the distributions to convey statistics. Median values for *v*_obs_ (m*_v_*) and PLCγ1-mNG density (m_ρ_) are expressly denoted for clarity. Lipid compositions for all experiments (mol%): 94:4:2 DOPC:Ni-DGS:PIP_2_. LAT densities: ∼1500 µm^-2^.

LAT condensates have the potential to exclude membrane proteins [8]. With membrane-tethered Hck as the primary source of PLCγ1 phosphorylation, we considered the possibility that pLAT condensation excludes Hck, leading to slower PLCγ1 phosphorylation and decreases in *v*_obs_ in condensates. To see if that was the case, we pre-phosphorylated PLCγ1 using BTK-KD in solution before addition to pLAT bilayers. Observed rates for pPLCγ1 were significantly higher than PLCγ1 phosphorylated by Hck alone, but the rate still dropped after introducing Grb2:SOS-PR (Fig 5C-D). If the m*_v_* for condensed pPLCγ1 were computed relative to uncondensed pPLCγ1 (rather than PLCγ1), it would be 0.39, equivalent to the relative rate for the experiment with no pre-phosphorylation (Fig 5B). Of note, the experiments in Fig 5A-B had labeled LAT-Ax555 (not shown) and had control lysate or Gads/SLP-76 lysate in addition to PLCγ1-mNG. The experiments in Fig 5C-D were done with unlabeled LAT and with only PLCγ1-mNG lysate, indicating little effect from these variables on the result.

## Discussion

Studies on PLCγ1 activation have been done for many years and in many formats [19–21,32]. Distinct to the assay introduced here is quantitative measurement of both PLCγ1 recruitment and activity together in real-time. In addition, proteins in the supported membrane assay interact in a physiologically relevant geometry, including the ability to form condensates, and their spatial organization on the membrane can be tracked. The use of unprocessed cell lysates also allows for incorporation of proteins that are difficult to purify and for the introduction of mutants with relatively low effort and short turnaround times. As demonstrated previously, this remains a powerful strategy for the study of signaling reactions at the membrane [24]. The data we report here agrees well with studies on the mechanisms of PLCγ1 activation in T Cells showing pLAT and pY783 are required for full activation (Figs 3B and 6A) [6,21,33].

**Fig 6.**
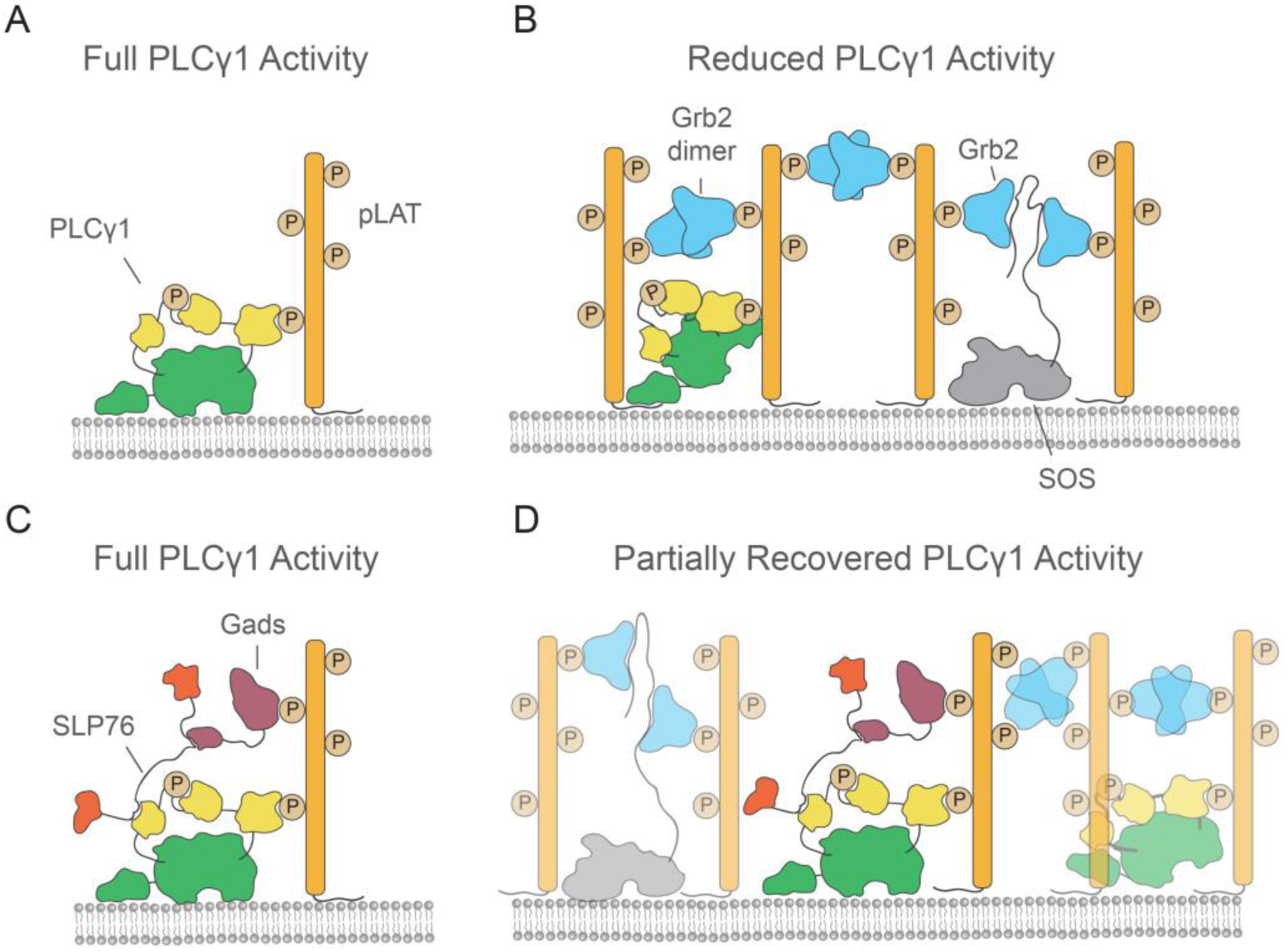
States of PLCγ1 activity in and out of the LAT condensate. (A) Graphic of PLCγ1 binding to pLAT and fully engaging the membrane, where full PLCγ1 activity is observed in experimental results. (B) A graphic of PLCγ1 in the LAT condensate, where PLCγ1 activity is reduced in experiments. Both Grb2 dimers and Grb2:SOS-PR interactions can crosslink LAT. (C) Graphic showing PLCγ1 interactions with Gads:SLP76 in the absence of a larger condensate. PLCγ1 is fully active in experiments with this configuration. (D) Graphic of PLCγ1:Gads:SLP76 interactions in the LAT condensate, where PLCγ1 activity is partially recovered compared to Grb2-only conditions.

Unique to the supported membrane assay is the ability to reveal how PLCγ1 function is affected by the condensation state of pLAT. Live cell research on membrane proximal T cell signaling has suggested essential elements of antigen discrimination occur between the recruitment of Zap70 and the activation of PLCγ1 [11,26,34,35]. This paints a clear target on pLAT and PLCγ1 as major players in signal gating from TCR to downstream calcium signals. PLCγ1 has already demonstrated the ability to partition into LAT condensates (and enhance their formation) in reconstituted experiments [23], and in live cells. Additionally, PLCγ1 activity in a parallel growth-receptor pathway condensation has been reported to increase [32]. Taken together, it would appear as if the incorporation of PLCγ1 into the LAT condensation would increase its activity, which would be a clear indicator of how LAT condensation could gatekeep T Cell activation.

However, our study shows pLAT condensation has a net *negative* effect on PLCγ1 activity relative to dispersed pLAT (Fig 5B). Simply enhancing enzyme concentrations or dwell time in a condensate is not always a positive regulator of signaling activity. This behavior of PLCγ1 stands in marked contrast to the Ras GEF SOS, which clearly depends on pLAT condensation as part of its activation process via a distinct type of kinetic proofreading [15,16,36], may be less obstructed due to its extended PR domain and differential binding via Grb2 (Fig 6B). The key observation presented here is that PLCγ1 is maximally active from dispersed pLAT. The observation of reduced activity in the condensate is likely system and reconstitution specific; we speculate that the density of the reconstituted condensate has the potential to sterically interfere with PLCγ1 access to its PIP_2_ substrate (Fig 6A-B). The inclusion of Gads:SLP76 in the condensate relieves some of the PLCγ1 inhibition, possibly by affecting the packing density around PLCγ1, but these effects were never observed to enhance activity relative to baseline conditions (Fig 6C-D). The supported membrane assay introduced here makes possible an increasing number of PLCγ1 studies and may facilitate the ultimate elucidation of how condensates regulate its signaling activity.

## Materials and Methods

### Cell transfection and lysate preparation

Full-length PLCγ1 with a C-terminal mNeonGreen (mNG) fusion protein (*mus musculus*), Gads (human), and SLP-76 (human) were cloned into a pCI backbone under the CMV promoter for expression in HEK293T cells (U47119.2).

HEK293T cells were transfected similarly to previously published methods [24]. HEK293T cells (UCB Cell Culture Facility) were seeded in clear bottom 6-well plates (Eppendorf) at a target confluency of 90% after allowing for adhesion overnight. Cells were cultured in DMEM medium, high glucose media (DMEM, Gibco) supplemented with 10% fetal bovine serum (heat shocked), minimal essential medium (MEM) non-essential amino acid and 100 μg ml−1 penicillin and streptomycin, and media was not changed before or after transfection. Cells were transiently transfected with exogenous vectors using Lipofectamine 3000 (Invitrogen) according to the manufacturer protocol for 6-well plates and with 3.75 µL volumes of the reagent. After allowing for >48 hrs of expression (no media change), transfection efficiency was evaluated using epifluorescence (for PLCγ1-mNG) and bright field microscopy. Typical efficiency (based on mNG fluorescence) was

>80%. After verifying high transfection efficiency, media was aspirated, and cells were collected in sterile 1X TBS. Cells were pelleted (1 well resulted in a ∼50 µL pellet) by gentle centrifugation and resuspended in 450 µL lysis buffer containing 20 mM Tris-HCl pH 7.4, 150 mM NaCl, 1% protease inhibitor cocktail (P8340, Sigma Aldrich), and 0.5% each of phosphatase inhibitor cocktails 2 (P0044, Sigma) and 3 (P5726, Sigma Aldrich).

Cells were lysed by 3 mm tapered tip sonication (Sonics VCX750 – 20% amplitude, 2 s sonication, 6 s rest, 20 cycles) in a 5 mL polypropylene tube immersed in an ice bath. Lysate was centrifuged at 20,000 g for 30 min at 4 °C. The supernatant was collected (leaving ∼50 µL in the tube), aliquoted, flash-frozen in liquid N_2_, and stored at -80 °C (used in experiments for up to ∼1 yr). Proteins seemed to maintain good stability (as demonstrated by PLCγ1 activity) without the use of glycerol, but 5-10% glycerol might be considered to increase stability.

### Western blots

1 µL of lysate per lane was thawed, treated as stated, and run on SurePAGE 4-12% Bis-Tris SDS–PAGE gels (GenScript) in 1X MES SDS-PAGE buffer and transferred to nitrocellulose membranes (Invitrogen) in 1X NuPAGE Transfer Buffer (Invitrogen). Approximate molecular weights were visualized using Precision Plus Protein Dual Xtra standards (Bio-Rad), which effectively transfer to membranes as well. Membranes were washed with 1X TBST and blocked with 5% BSA in TBST. The membranes were incubated with either PLCγ1 (D9H10) XP rabbit mAb (Cell Signaling Technology, 1:1000 dilution), phospho-PLCγ1 (Tyr783) (D6M9S) rabbit mAb (Cell Signaling Technology, 1:1000 dilution), SLP-76 (D1R1A) rabbit mAb (Cell Signaling Technology, 1:1000 dilution), or Gads (UW40) mouse mAb conjugated with Alexa 647 (Santa Cruz Biotechnology, 1:200 dilution) overnight at 4 °C with agitation. Membranes were then incubated with Goat anti-Rabbit IgG (H+L) conjugated to Alexa 647 (A-21244, Invitrogen, 1:20,000 dilution) at room temperature for 1 hr with agitation. Bands were visualized by fluorescence at 640 nm using a Typhoon FLA 9500 laser scanner (Cytiva). Band intensities and backgrounds were quantified using the Fiji software package.

### Solution kinase reactions

1 µL of lysate per reaction was thawed then diluted in reaction buffer containing 20 mM Tris-HCl pH 7.4, 150 mM NaCl, 20 mM MgCl_2_, 1 mM ATP (Sigma Aldrich), 1 mM activated sodium orthovanadate (508605, Calbiochem). Either Milli-Q water, 10 µM His6-Hck, or 10 µM BTK-KD was added for a final volume of 12 µL. After the stated time, reactions were quenched with 3 µL of 5X SDS-PAGE loading buffer. The entire 15 µL volume was loaded for SDS-PAGE, and results were visualized by western blot as above.

### Protein purifications

All purified constructs (save for BTK-KD, see below) utilized pET28a (Novagen-Cat. # 69864-3) or pETM (similar to Addgene #108943) backbones under the T7 promoter for expression in *E. Coli*. Expression vectors were transformed into BL21-CodonPlus competent *E. Coli* (Agilent) and cultured in TB. T7 expression was induced using 2 mM IPTG (final concentration) (Gold Biotechnology) overnight (18-20 hrs) at 18 °C with shaking. Pelleted cells were resuspended in bacterial lysis buffer typically containing 75 mM NaH_2_PO_4_ pH 8.0, 300-700 mM NaCl, 2 mM beta-mercaptoethanol (Sigma Aldrich), DNAse (Sigma Aldrich), 5-10 mM imidazole (Fisher Scientific), and EDTA-free Pierce Protease Inhibitor Cocktail (A32965, ThermoFisher Scientific). All subsequent steps were done at 4 °C. Bacteria were lysed using an EmulsiFlex-C3 high-pressure homogenizer (Avestin). Subsequent steps are described for each protein below, with proteins generally stored at -80 °C in 20 mM Tris-HCl (pH 8 at 4 °C), 200 mM NaCl, 1 mM TCEP (Sigma Aldrich), and 10% glycerol (MCB grade, Fisher Scientific) after a final SEC polishing step (Superdex 75 or 200 column, 120 mL bed volume, Cytiva) and flash-freezing in liquid N_2_.

Full-length Grb2 (human) was expressed with a cleavable (TEV) N-terminal His6-tag and purified as described previously [16]. Grb2 was cleaved with TEV to remove the His6-tag. Of special note, Grb2 was incubated at 37 °C for 15 min in a thermocycler to allow dimerization before using it in experiments [16].

PKCθ-C1b domain (232-282 aa, *mus musculus*) (“DAG Sensor”) was expressed with a C-terminal SNAP-tag fusion protein (SAGGSASAGGSA linker) and purified equivalently to Grb2 using an N-terminal His6-tag. Of special note, growth media for C1b expression was supplemented with 0.01 mM ZnSO4 (final concentration) [37]. PKCθ-C1b-SNAP was cleaved with TEV to remove the His6-tag.

LAT (30-233 aa, human) was expressed with an uncleavable His10-tag with a SAGGSASAGGSA linker and with the mutations: Y36F, Y45F, Y64F, Y70F, and Y110F. LAT was otherwise purified as described previously [38].

Hck (83-526 aa, human) was expressed with a cleavable (TEV) His6-tag and purified as described previously for Src-family kinases [39]. The His6-tag was left intact for membrane tethering. Of special note, the purification strategy utilized calls for co-expression of YopH phosphatase in the *E. Coli* host, which is essential to separate out for pLAT experiments. YopH seems to bind extensively to Hck (while running at a similar molecular weight in SDS-PAGE) and should be removed by high salt washes and careful ion exchange chromatography.

SOS-PR domain (1,051–1,333 aa, human) was expressed with a cleavable (TEV) N-terminal His6-MBP-fusion tag and purified as described previously [10]. Of special note, it is essential that this construct be purified by an ion exchange step (MonoS column, 1 mL bed volume, Cytiva) with high amounts of reducing agent (>10 mM BME for example).

SOS-PR seems to form strong adducts with MBP and bacterial contaminants which can strongly affect results downstream, but these adducts can be disrupted by ionic differences and reducing agents. SOS-PR was cleaved with TEV to remove the His6-MBP-tag before use.

BTK kinase domain (402-659 aa, human) was expressed in insect cells and purified as described previously for other kinase domains [38].

PLCδ pleckstrin homology domain (11-140 aa, human) (“PIP_2_ Sensor”) was purified and labeled as previously described [27].

### Fluorescent labeling of proteins

PKCθ-C1b-SNAP (DAG sensor) was labeled with SNAP-Surface Alexa 647 (SNAP-Alexa 647, S9136S, New England Biolabs) according to the manufacturer protocol. LAT was labeled at residue C117 using Alexa 555 C2 maleimide (Ax555, Invitrogen) according to the manufacturer protocol. Briefly, LAT was diluted to 100 µM with 5 mM TCEP, then allowed to react with the maleimide dye for 2 hrs at room temperature before quenching with 10 mM BME. Excess dye was initially removed by gravity desalting columns (PD10, Cytiva) and further by SEC via FPLC (Superdex 75 increase, 24 mL bed volume, Cytiva). Of special note, we have observed LAT to appear as multiple SEC peaks (up to 3 or more), but these do not seem to be distinct populations for our purposes (if isolated, they all behave nominally the same on SLBs). We suggest simply pooling all SEC peaks of LAT (given they are sufficiently pure). Labeled fraction was determined by comparison of absorption at the dye’s λ_max_ to the protein’s UV absorption at 280 nm (after correction for the dye’s contribution at 280 nm, according to manufacturer protocol).

### SUV and SLB preparation

Small unilamellar vesicles (SUVs) were made using 1,2-dioleoyl-sn-glycero-3-phosphocholine (DOPC), 1,2-dioleoyl-sn-glycero-3-[(N-carboxypentyl)iminidiacetic acid)succinyl] (nickel salt) (Ni2+-NTA-DGS), and L-α-phosphatidylinositol-4,5-bisphosphate (Brain, Porcine) (ammonium salt) (Brain PIP_2_) (Avanti Polar Lipids). Lipids were purchased, stored (-20 °C), and mixed solubilized in chloroform. Lipids were mixed in an etched round bottom flask to yield 2 mg total lipid mass at a ratio of 94:4:2 (mol%) DOPC:Ni-DGS:PIP_2_. After mixing well, lipids were dried by rotovap in a 37 °C water bath, kept under vacuum to dry an additional 10 min at room temperature, then dried under continuous N_2_ gas flow for >30 min. Lipid films were solubilized with 2 mL milli-Q water (to yield 1 mg/mL lipid concentration) and vigorously vortexed to ensure film liftoff. The cloudy aqueous lipid mixture was added to a 5 mL polypropylene tube and treated by 3 mm stepped tip sonication (Sonics VCX750 – 33% amplitude, 20 s sonication, 50 s rest, 5 cycles) while submerged in ice. The clarified SUV solution was then centrifuged at 20,000 g for 20 min to remove tip debris and any remaining large vesicles. The supernatant was stored at 4 °C in a sealed tube for use in experiments up to ∼1 week.

Precision Schott D 263 M glass coverslips (#1.5H, 75 x 25 mm, custom ordered from ThorLabs, similar to Part# CG15CH) were first cleaned with warm 2% Hellmanex III (Fisher Scientific) solution with bath sonication at 45 °C for 30 min, then 50% IPA:mill-Q with sonication for >15 min, then milli-Q with sonication for >15 min. Substrates were then etched using a 1:3 ratio of concentrated H_2_SO_4_ to 30% H_2_O_2_ (piranha etch) for 3 min (while still warm). Substrates were rinsed excessively with milli-Q water between each cleaning step and after etching. Substrates were stored in milli-Q at room temperature for use in experiments up to ∼1 week. To build chambers for experiments, etched substrates were blow-dried under N_2_ and attached to simple 6-channel flow cells (sticky-Slide VI 0.4, ibidi). Chambers were cured at 37 °C for >10 min while pressed under a heavy weight. After curing, chambers were allowed to cool to room temperature for >10 min.

Supported lipid bilayers were formed via vesicle rupture. The 1 mg/mL SUV solution was diluted to 0.25 mg/mL in degassed 1X TBS. 100 µL of 0.25 mg/mL SUV (0.75X TBS) was added per chamber channel and incubated for >30 min at room temperature. Channels were washed via 1 mL degassed 1X TBS. SLBs were then blocked with 1 mg/mL casein solution (Blocker Casein, Thermo Scientific) diluted in 1X TBS for 10 min. Channels were exchanged to an incubation buffer (1X TBS, 2 mM MgCl_2_, 1 mM TCEP). His10-LAT and His6-Hck were thawed, diluted in incubation buffer, then centrifuged at 20,000 g at 4 °C to remove aggregates before use. 100 nM His10-LAT and 10 nM His6-Hck in incubation buffer (with 0.1 mg/mL casein) were added to bilayers (100 µL per channel) and incubated for 20 min (yields ∼1500 µm^-2^ LAT density). After an initial wash, channels were incubated in 1 mM ATP, 1 mM sodium orthovanadate, and 0.1 mg/mL casein in incubation buffer for 20 min to both phosphorylate LAT via Hck and to allow weakly chelated proteins to dissociate. After a secondary wash, channels were ready for experiments (used within ∼1-4 hrs).

### Imaging Hardware and Acquisition

TIRF imaging experiments were performed on an inverted Nikon Eclipse Ti microscope using a 100X Nikon oil-immersion TIRF objective (1.49 NA). We used a 1.5X lens tube (Nikon) before the camera for a total 150X magnification. Micro-positioning was done using a motor-controlled stage (MS-2000, ASI). Images were acquired with an iXon Ultra 897 EMCCD camera (Andor Technology). Fluorescently labeled proteins were excited with either a 488-, 561-, or 637-nm diode laser (OBIS laser diode; Coherent) controlled with a custom laser driver (Solamere Technology) using TTL signals from the camera. Lasers were collimated with the objective position at the approximate sample z-focus before every experiment. A single quad-band dichroic cube was used for all acquisitions (ZT405/488/561/640rpcv2, Chroma). Fluorescence emission was additionally filtered by: ET525/50M (488 excitation), ET600/50M (561 excitation), or ET700/75M (637 excitation) (Semrock). Microscope and accessory hardware were controlled using Micro-Manager, version 4.0 [40]. Experiments were conducted at 21-23 °C. Excitation conditions for each channel during background collection and kinetic measurements were as follows: PLCγ1-mNG was excited at 488 nm (∼0.45 mW at the objective), LAT-Ax555 was excited at 561 nm (∼0.25 mW at the objective), and DAG sensor was excited at 637 nm (∼0.2 mW at the objective). All channels were acquired with a 20 ms camera exposure and 500 EM gain, with 5-10 s delays between time points for time-lapse measurements. Images for kinetic traces were taken with the objective focused on the center of the channel of interest. Background images (n = 20-30) were collected before and after kinetic measurement to be used for field illumination (“shade”) correction (to remove aberrations and uneven TIRF illumination). Corrections and quantitative image analyses were done in MATLAB.

### Lysate assay kinetic acquisitions

All kinetic measurements were performed using imaging buffer containing 1X TBS, 2 mM MgCl_2_, 1 mM TCEP, 1 mM ATP, 1 mM sodium orthovanadate, 2 mM Trolox (UV treated), 10 mM BME, 0.1 mg/mL casein (final concentrations after addition of analytes/sensors). Trolox (Sigma Aldrich) was prepared as previously described [27]. 1 µL of lysate and 200 nM DAG sensor were used unless otherwise stated. When adaptors were included (Grb2/SOS/Gads/SLP-76), components were combined at stated concentrations and added as a single injection. DAG sensor was first incubated in channels for >10 min in imaging buffer to collect background images, allow nonspecific equilibration of the sensor at the membrane, and aid in finding the objective z-focus. Background measurements were taken for 30 s before carefully injecting lysate/DAG sensor solutions mid-acquisition (such that the objective focus was unaffected). Images were collected at 5-10 s intervals for as long as 1.5 hrs.

### Image Analysis and Segmentation

Quick visualizations and quality checks of TIRF images were done using FIJI [41]. For quantitative analysis, images were imported into MATLAB using custom scripts. MATLAB was then used for cropping, field correction, patch segmentation, and subsequent trace analyses. For kinetic traces, the raw value for each timepoint is the median pixel intensity of the corresponding patch. Traces were then normalized to the time of injection (for each patch, separately). Calibrations were then applied as described below. For the creation of the LAT condensation mask (Fig 4D), the built-in MATLAB function imgaussfilt was used (σ = 1) and an empirically determined threshold equivalent to the median (filtered) intensity was applied. All plotting was also done using MATLAB.

### Quantitative calibrations

LAT-Ax555 density was calibrated as previously described [10]. Briefly, a series of SLBs with increasing LAT density were prepared. TIRF images were taken of each channel, followed immediately by Fluorescence Correlation Spectroscopy (FCS) measurements. Densities calculated from FCS fits were correlated with median TIRF intensities to create a calibration curve.

For the conversion of raw fluorescence to DAG (in mol%), normalized DAG sensor traces were first cataloged for several experiments (with 1 µL PLCγ1-mNG lysate) of over multiple days (n = 10). Given that all experiments had approximately 2 mol% PIP_2_ in the bilayer (with some random error due to imperfect mixing, etc.), it was assumed the DAG sensor trace should plateau at 2 mol% DAG after full conversion of PIP_2_ (which was typically close to 20 min of reaction time for this condition). The median for all cataloged traces (𝐼_𝑛𝑜𝑟𝑚. 𝑚𝑒𝑑._) was computed, and an “intensity to DAG” coefficient, 𝜀_𝐷𝐴𝐺_, was defined and computed such that % 𝐷𝐴𝐺 = 𝐼_𝑛𝑜𝑟𝑚. 𝑚𝑒𝑑._ × 𝜀_𝐷𝐴𝐺_. The empirical value for 𝜀_𝐷𝐴𝐺_ was calculated to be 0.880±0.08 (mol% DAG) / (norm. intensity) (1σ, n = 10), indicating an acceptable consistency in the measurement.

For PLCγ1-mNG density (in µm^-2^), TIRF images of PLCγ1-mNG were taken at varying concentrations that allowed for single molecule counting (at high laser powers). Images at the low laser power used for kinetic measurements (∼0.45 mW at the objective) and images at high laser power (∼10 mW at the objective) were taken in quick succession. The high-power images were analyzed using the TrackMate plugin of FIJI in order to count the number of PLCγ1-mNG present [42]. The single molecule counts were then correlated with the low-power images to calculate the approximate density of PLCγ1-mNG from their TIRF fluorescence intensity (background corrected).

### *v*_obs_ measurements

Given that PLCγ1 activation is likely a multistep process, the true enzymatic rate cannot be measured by bulk measurements (the measured PLCγ1-mNG density is not necessarily the active density). Instead, we defined an observed velocity, *v*_obs_, to be the linear steady-state rate of DAG production apparent in our data and utilized this for comparison of varied experimental configurations. To calculate *v*_obs_, the linear portion for every DAG curve (patch by patch) was determined by finding the best linear fit (highest r^2^) for a moving time-axis window. In detail: for every curve, in MATLAB, we performed a scan from *t*_0.1_ (time to reach 10% of the maximum) to *t*_0.9_ (time to reach 90% of the maximum). Through linear regression, we calculated the slope and r^2^ correlation coefficient using a moving fixed-size time window across this section. The slope corresponding to the highest r^2^ was stored as the *v*_obs_ for the curve in question.

## Acknowledgments

We thank the members of the Groves Laboratory for helpful discussion and critical feedback.

## Notes

### Competing Interest Statement

The authors have declared no competing interest.

### Funding

- Novo Nordisk Foundation Challenge Programme: Center for Geometrically Engineered Cellular Systems NNF17OC0028176
- NIH NIAID: P01 AI091580

