## Supplementary Information for "Supported membrane assay probes PLCγ1 activity in LAT condensates"

*Corresponding Author

**This PDF file includes:**

Figures S1 to S3

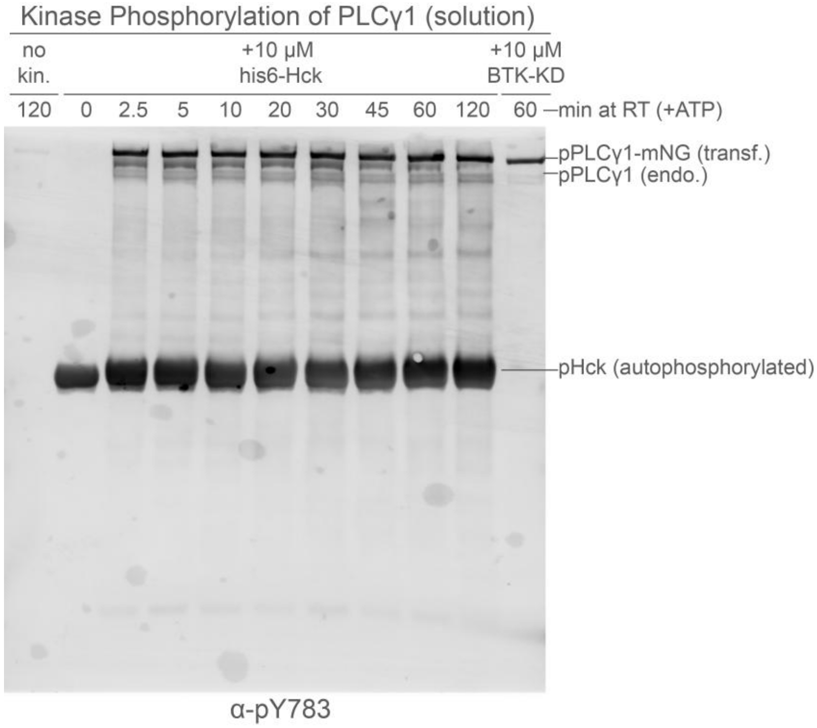


**S1 Fig. Full Western Blot of in vitro PLCγ1 phosphorylation.** The full Western Blot from Figure 2C. Western blot probed for pY783 with a series of time points and different kinase conditions. Phosphorylation reactions were conducted in 10 μL volumes containing 1 μL of PLCγ1-mNG lysate each, kept at RT for the designated time, then quenched with SDS-PAGE loading buffer. Either no kinase, His6-Hck (FL), or BTK-KD (Kinase Domain) was used. While α-PLCγ1-pY783 antibody was used, other phosphoproteins were also detected in the background. pHck appears prevalently as a large band in lanes where it was included. Hck lanes also demonstrate a much higher general background, with many proteins in the lysate presumably being phosphorylated. In contrast, BTK-KD shows more specific activity for PLCγ1-Y783.

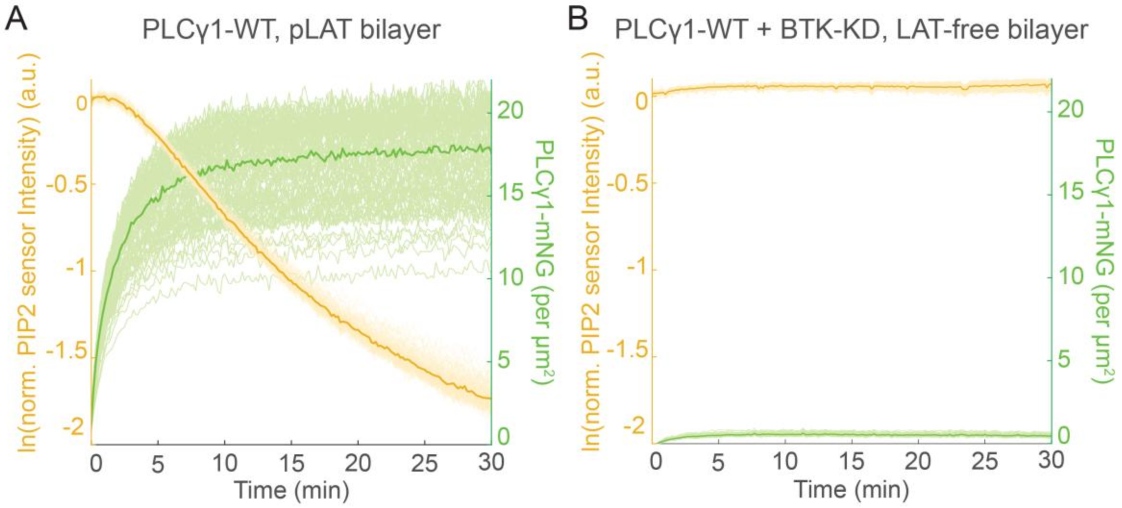


**S2 Fig. Lysate assay with PIP2 sensor and pPLCγ1-mNG on a LAT-free bilayer.** (A-B) Curves obtained from processing the raw median fluorescence from membrane patches, representing the amount of PIP2 in the bilayer (gold, left axes, top curves) and density of PLCγ1-mNG recruited (green, right axes, bottom curves) over 30 minutes for either a pLAT functionalized bilayer (A) or a LAT-free bilayer with pre-phosphorylated pPLCγ1-mNG (B). pPLCγ1-mNG was generated by incubating PLCγ1-mNG with 1 μM BTK-KD for 1 hr prior to injection. PLCγ1-mNG lysate (~1:1000 dilution) with PIP2 sensor (Cy3-PLCδ-PH) was injected at t = 0 min. A decay in PIP2 sensor intensity correlates to hydrolysis by PLCγ1-mNG. PLCγ1-mNG density curves were attained using calibrations after normalization to the time of injection. Solid, darkly colored curves represent the median of all membrane patches analyzed. Lightly colored curves in the background represent the results from each single patch (n = 100).

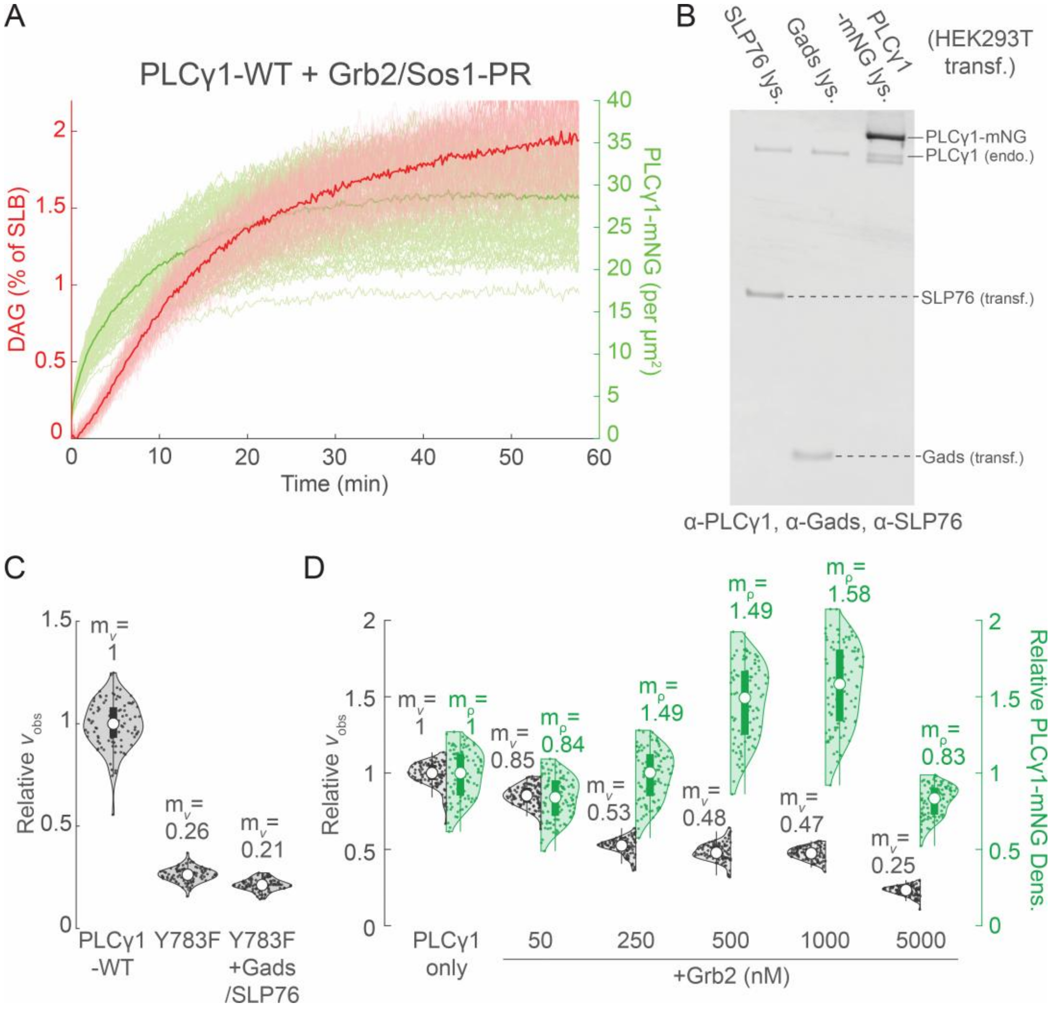


**S3 Fig. Validation of the lysate assay with the addition of the LAT condensate.** (A) Curves obtained from processing the raw median fluorescence from membrane patches, representing the amount of DAG in the bilayer (red, left axis, top curves) and density of PLCγ1-mNG recruited (green, right axis, bottom curves) over 60 minutes. The production of DAG slows considerably but eventually reaches the maximum possible 2% of the SLB. (B) Western blot demonstrating the expression of Gads-FL and SLP76-FL in HEK293T cells. Gads and SLP76 containing lysates were used for (C) and Figure 5A-B. (C) Relative $v_{obs}$ for PLCγ1-WT (with 2 μL of control lysate), for PLCγ1-Y783F mutant (with 2 μL of control lysate), and PLCγ1-Y783F mutant + Gads/SLP76 (1 μL lysate each). Data were normalized to the median value of PLCγ1-WT. There is not a demonstrative difference between the lysate batches, nor does the addition of Gads/SLP76 result in a strong activation of Y783F mutant. (D) Relative $v_{obs}$ and relative PLCγ1-mNG densities for various conditions, with an increasing amount of purified Grb2. Data were normalized to the median value of the PLCγ1-only condition. Grb2 has been shown to have the ability to crosslink LAT on its own due to dimerization.
